# Can Metabolic Thresholds be used as Exercise Intensity Markers in Adult men with Obesity?

**DOI:** 10.1101/440958

**Authors:** Peric Ratko, Nikolovski Zoran

## Abstract

The first aim of the study was to identify the exercise intensity eliciting the highest (FAT_max_) and the lowest (FAT_min_) fat oxidation rate in men with obesity. The second aim was to evaluate if FAT_max_ and FAT_min_ correlate with aerobic (AeT) and anaerobic (AnT) thresholds, which in turn could be used as exercise intensity markers. Nineteen adult sedentary men participated in the study. Breath-by-breath analysis was performed throughout the test to assess maximal oxygen consumption (VO_2max_) with stoichiometric equations used to calculate fat oxidation rate. Pearson correlation coefficient (*r*), coefficient of determination (*R^2^*) and paired t-test were used to evaluate differences between VO_2_ at AeT and at FAT_max_ and VO_2_ at AnT and at FAT_min_, respectively. FAT_max_ and AeT occurred at 42.80 ± 2.68 % of VO_2max_ and 43.02 ± 2.73 % of VO_2max_, while FAT_min_ and AnT occurred at 53.40 ± 3.65 % of VO_2max_ and 53.38 ± 3.65 % of VO_2max_, respectively. A high correlations were found between VO_2_ at FAT_max_ and at AeT (*r* = 0.86, *p* < 0.01) and VO_2_ at FAT_min_ and at AnT (*r* = 0.99, *p* < 0.01). The existing correlations suggest that metabolic thresholds may be used as exercise intensity markers in men with obesity.

## Introduction

The current obesity epidemic is considered one of the leading causes of various conditions and chronic diseases worldwide [1,2]. Over the last decades, studies demonstrated that obesity-associated conditions such as type 2 diabetes, cardio-vascular diseases and certain types of cancer are the primary causes of premature mortality [2,3].

In addition to lifestyle and diet changes, physical activity via aerobic exercise can directly influence obesity-associated conditions by improving weight control and the metabolic rate [4,5]. Aerobic exercise at the intensity eliciting a maximal rate of fat oxidation (FAT_max_) revealed best results in treatment and prevention of obesity-associated conditions by means of increase in maximal fat oxidation rates (MFO) and insulin sensitivity, demonstrating a significant role in the treatment of obesity-associated conditions [2,3]. As a follow-up, several studies demonstrated how the lowest beneficial effect was observed when aerobic exercise was performed at intensity matching negligible fat oxidation rate (FAT_min_), contributing to diminished protection of exercise therapy [6–8]. Therefore, choosing appropriate exercise intensity may play a decisive role in decreasing risk factors accompanying obesity-associated conditions.

Exercise intensity prescription is commonly expressed as a percentage of maximal oxygen consumption (% VO_2max_) [2]. However, this method usually leads to a existence of high inter-subjects variability and wide “FAT_max_ zone” (range 30 % to 65 % of VO_2max_), compromising the accuracy of exercise prescription in men with obesity [4,9,10]. Although the relative percentage concept is widely used, recent criticism has emerged concerning its precision, and usage of heart rate (HR) corresponding to the metabolic thresholds as reference for more individual exercise prescription has been advocated [1–3].

Obtained during a graded exercise test (GXT), either noninvasively via ventilatory gases (VT) or invasively by determination of blood lactate concentration (LT), first metabolic threshold (aerobic (AeT)) and second metabolic threshold (anaerobic (AnT)) are individual indicators of exercise intensity and can be used to accurately delineate intensity zones for more efficient aerobic exercise prescription [11,12].

As fat oxidation is directly dependent on the availability of oxygen (O_2_), the intensity that elicits the FAT_max_ and FAT_min_ might be related to the intensity at the AeT and AnT. Therefore, the prescription of aerobic exercise based on the HR at the metabolic thresholds may be a valid strategy for more specific exercise therapy in individuals with obesity moving away from the traditional “FAT_max_ zone” concept.

With intention of improving the effectiveness on exercise prescription, studies examined the connection between the metabolic thresholds and fat oxidation in various subjects [7,8,13]. However, only low to modest correlations between FAT_max_ and AeT have been reported in men with obesity during a cycle ergometry with no available data regarding treadmill testing [9,14]. Furthermore, high correlations between FAT_min_ and AnT have been observed in sedentary and athletic population with no studies available regarding obese [7,16]. Consequently, the relationship between metabolic thresholds and the fat oxidation rate is still not clear in sedentary adult men with obesity during their natural movement, walking and running, directly reducing efficiency of exercise prescription on health promotion in obesity-associated conditions.

Therefore, the first aim of this study is to determine FAT_max_ and FAT_min_ during GXT performed on a motorized treadmill. The second aim is to examine the correlation between VO_2_ at FAT_max_ and at AeT and between VO_2_ at FAT_min_ and at AnT when thresholds are assessed via ventilatory gases in adult sedentary men with obesity. We hypothesized existence of strong correlation between VO_2_ at FAT_max_ and at AeT and between VO_2_ at FAT_min_ and at AnT, which in turn may be used as exercise intensity marker for more individual training prescription in observed population.

## Materials and methods

### Subjects

Nineteen adult sedentary men with obesity completed the study. All subjects were classified as obese if body fat ≥ 30 % [16]. They were recruited from circulated flyers and advertisements in the newsletters. All participants were familiarized with ethical principal of the study, potential health risks and objectives and willingly singed consent statement regarding publication of the data. Entire research complies with the policy statement with respect to the Declaration of Helsinki and was approved by the Institutional ethical committee. Prior to testing, all participants completed a questionnaire regarding their health status and activity. All subjects were nonsmokers and presented no evidence of health disorders. Participants were excluded from the study if diabetic, hypertensive (≥ 140/90□mmHg), if practiced any organized physical activity prior to this study or if they were consuming medication which might affect substrates utilization. Subjects were requested to abstain from any physical activity 48 h prior to the testing. Also, they were asked to refrain from consuming caffeine and nutritional supplements on the testing day. Anthropometric characteristics and bioimpedance analysis (BIA) were assessed before testing at the first laboratory visit using a combined stadiometer and a BC - 418 Segmental Body Composition Analyzer (TANITA_®_ Tokyo, Japan). Testing was performed between 08:00 and 12:00 h in a laboratory with standardized environmental conditions (21 °C, 43% humidity and 270 m above sea level). The prescription of low on carbohydrate (CHO), high on protein and fat (LCHPF) diet is considered as possible strategy for weight loss and as such preferred treatment for obesity [17]. Therefore, last pretest LCHPF meal (15 % CHO, 55 % protein, 40 % fat), not representative of the subject’s habitual diet, was consumed 12 h before test, with test sessions being performed in a fasted state [18]. To prevent excretion of (non-oxidative) carbon dioxide (CO_2_), which can overestimate CHO and underestimate fat oxidation at high intensities, subjects whose maximal breathing frequency (BF_max_) during test was ≥ 45 breaths per minute (breaths·min^−1^), were excluded from the study [19,20].

### Exercise protocols

Participants performed GXT on a T170D motorized treadmill (COSMED_®_, Rome, Italy) until volitional exhaustion to measure VO_2max_ with substrates utilization being measured continuously. Oxygen consumption (VO_2_) and carbon dioxide production (VCO_2_) were assessed during test by breathe-by-breath Quark PFT Ergo (COSMED_®_, Rome, Italy) system. The flow sensor and the gas analyzers were calibrated using a 3-L syringe and calibration gas (O_2_ 16.10 % and 20.93 %; CO_2_ 0.00 % and 5.20 %) before each test. Fluctuations of breath-by-breath data were minimized using a 6 breaths smoothing and consequent 30 sec averaging as suggested by the manufacturer. Metabolic thresholds were determined manually via ventilatory gases using following criteria: the first rise in the ventilatory equivalent for O_2_ without concomitant rise of ventilatory equivalent for CO_2_ for AeT and V-slope method for AnT [11,21]. VO_2max_ was determined using the following criteria: a respiratory exchange ratio (RER) ≥ 1.15 or a plateau of VO_2_ in spite of a load increase; measured in liters per minute (L·min^−1^), normalized and expressed in milliliters per kilogram of body weight per minute (ml·kg^−1^·min^−1^) [22]. Participant’s HR was monitored continuously with integrated monitoring system (POLAR_®_, Kempele, Finland) and presented in beats per minute (beats·min^−1^). The GXT protocol used consisted of three stages: rest, exercise and recovery. The resting stage (3 min) was performed in seating to obtain baseline values and assure low starting HR. The exercise stage initial speed was 4 km·h^−1^ followed by an increase in speed of 1 km·h^−1^ every 2 min until volitional exhaustion. Incline was kept constant at 1 %. The subjects were instructed to self-select when the transition from a walk to a run occurs. Exercise stage was followed by a 3 min active recovery at the 4 km·h^−1^ in order to observe physiological recovery of the subjects. All participants underwent 15 min walking familiarization a day before test to get used to the breathing apparatus and treadmill.

### Substrates calculation

Average values for VO_2_ and VCO_2_ were calculated throughout exercise and collected at mouth level. For each subject, the values of substrates utilization were calculated from revised stoichiometric equations with the assumption that the urinary nitrogen excretion was negligible [23,24]. Obtained data were depicted graphically as a function of fat oxidation versus exercise intensity expressed in VO_2_. The resulting graphs were then used to determine the exercise intensity at which the FAT_max_ and FAT_min_ occurred, as well as the absolute (g·min^−1^) and relative (%) fat oxidation rates for each individual.

### Statistics

All data are analyzed and presented as mean ± standard deviation (*SD*), 95 % confidence intervals (*CI*) and range using the MedCalc (MEDCALC_®_, Ostend, Belgium) software. Normal distribution of all measured variables was addressed using the *Shapiro-Wilk* test with normal distribution accepted for all variables. The strength of the relationship between VO_2_ at AeT and VO_2_ at FAT_max_ was assessed using the Pearson product moment (*r*) correlation coefficient with same being done for AnT and FAT_min_ correlation. A coefficient of determination (*R^2^*) was used to detect the proportion of existing variance. A paired *t*-test was used to asses’ differences between measured variables. For all statistical analyses, significance was accepted at *p* ≤ 0.05.

## Results

All participants successfully finalized the study. Table 1 summarizes participants’ anthropometric characteristics. Observed mean VO_2max_ was 40.26 ± 2.75 ml·kg^−1^·min^−1^ (range 35.65 to 45.89; 95 % CI 38.93 to 41.58). Mean maximal HR (HR_max_) was 169.21 ± 5.76 beats·min^−1^ (range 161 to 180; 95 % CI 166.43 to 171.99). Mean BF_max_ was 38.1 ± 5.2 breaths·min^−1^ (range 34 to 44; 95 % CI 37.23 to 42.56).

**Table 1.** Participants’ anthropometric characteristics (*n* = 19)

| Variable | Mean ± SD |
| --- | --- |
| Age (y) | 32 ± 6.7 |
| Body mass (kg) | 121.25 ± 6.61 |
| Height (cm) | 175.25 ± 4.49 |
| Body fat (%) | 34.75 ± 2.60 |

\* *n* number of participants

Participants reached AeT at 17.32 ± 1.10 ml·kg^−1^·min^−1^ of VO_2_ (range 15.69 to 19.25; 95 % CI 16.78 to 18.85) or 43.02 ± 2.73 % of VO_2max_ (range 38.97 to 47.81), respectively. Mean HR at AeT was 109 ± 2 beats·min^−1^ (range 105 to 115; 95 % CI 107 to 110) or 66.46 % HR_max_. FAT_max_ was reached at a 17.23 ± 1.02 ml·kg^−1^·min^−1^ of VO_2_ (range 15.69 to 18.96; 95 % CI 16.73 to 17.72) or 42.80 ± 2.68 % of VO2max (range 38.97 to 47.09).

AnT was obtained at 21.49 ± 1.47 ml·kg^−1^·min^−1^ of VO_2_ (range 19.89 to 25.17; 95 % CI 20.79 to 22.20) or at 53.38 ± 3.65 % of VO_2max_ (range 49.40 to 62.52), respectively. Mean HR at AnT was 141 ± 3 beats·min^−1^ (range 137 to 148; 95 % CI 139 to 143) or 85.98 % HR_max_. FAT_min_ was achieved at 21.50 ± 1.47 ml·kg^−1^·min^−1^ of VO_2_ (range 19.89 to 25.17; 95 % CI 20.79 to 22.21) or at 53.40 ± 3.65 % of VO_2max_ (range 49.40 to 62.52).

The relative MFO at FAT_max_ was 76.42 ± 16.06 % of total energy share (95 % CI 66.21 to 86.62; range 45.00 to 96.00). Our participants absolute MFO was 0.72 ± 0.31 g·min^−1^ at FAT_max_ (95 % CI 0.52 to 0.92; range 0.20 to 1.12). The relative MFO at FAT_min_ was 0.00 ± 0.00 % of total energy share (95 % CI 00.00 to 00.00; range 00.00 to 00.00). The absolute MFO at FAT_min_ was 0.01 ± 0.01 g·min^−1^ (95 % CI - 0.01 to 0.01; range 0.00 to 0.01).

A Pearson correlation coefficient revealed a high correlation between VO_2_ at AeT and at FAT_max_ (*r* = 0.89, *p* < 0.01, 95 % CI 0.72 to 0.96) (Fig 1). The AeT and FAT_max_ connection showed large coefficient of determination (*R^2^* = 0.7921) accounting for 79.21 % of the variance. A correlation coefficient analysis between VO_2_ at AnT and at FAT_min_ (*r* = 0.99, *p* < 0.01, 95 % CI 0.99 to 1.00) was high (Fig 2). The AnT and FAT_min_ connection exhibited large coefficient of determination (*R*^2^ = 0.9801) accounting for 98.01 % of the variance, respectively.

**Fig. 1.**
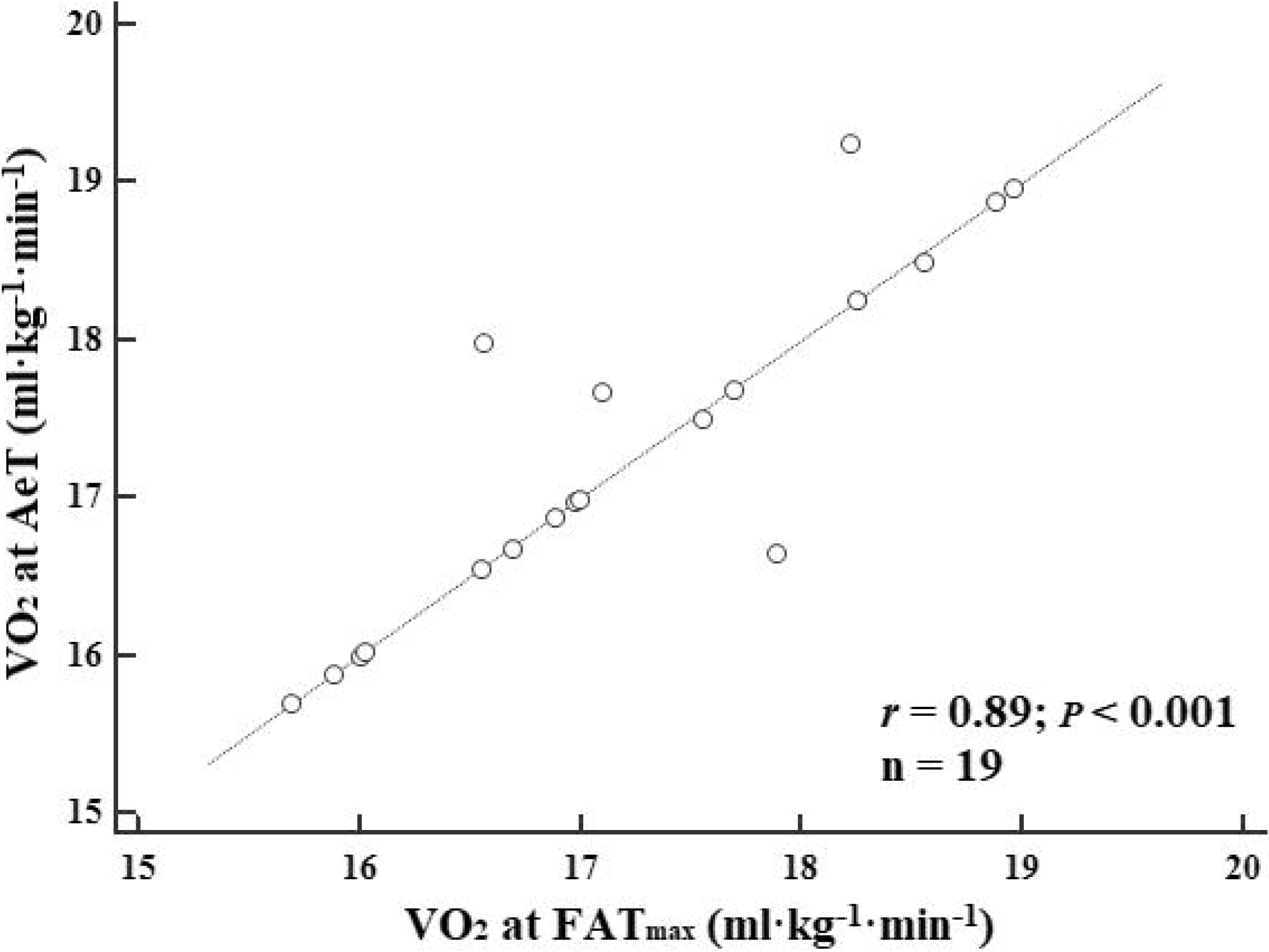
A Pearson correlation coefficient between VO_2_ at AeT and VO_2_ at FAT_max_ with line of equality (*n* = number of subjects)

**Fig. 2.**
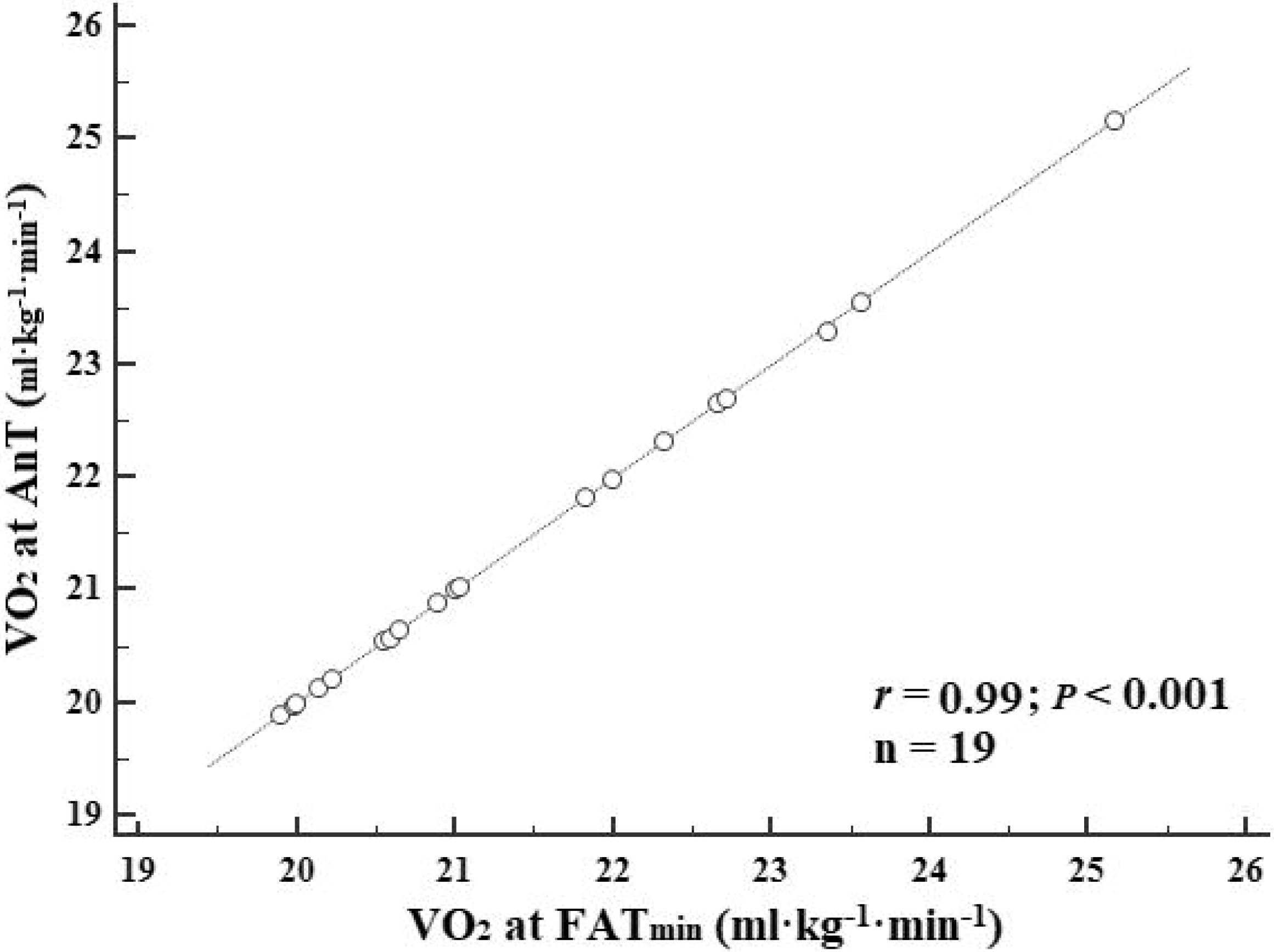
A Pearson correlation coefficient between VO_2_ at AnT and VO_2_ at FAT_min_ with line of equality (*n* = number of subjects)

A paired t-test demonstrated no statistical difference betwen VO_2_ at FAT_max_ and at AeT (*t*-value = 0.75, *p* = 0.46). The same was observed for the VO_2_ at FAT_min_ and at AnT (*t*-value = 1. 51, *p* = 0.15).

## Discussion

This was the first study to determine the exercise intensities eliciting the FAT_max_ and FAT_min_ in adult sedentary men with obesity using a short stage GXT treadmill protocol and to successfully correlate these intensities with their individual AeT and AnT when assessed via ventilatory gases.

In the present study, FAT_max_ occurred at intensity of 42.80 % VO_2max_ (range 38.97 to 47.09) which is in line with previous investigations [4,10,14]. These results confirm how the fat oxidation is optimal at low-to-moderate exercise intensities in subjects with obesity [7]. Nevertheless, direct comparisons with previous studies are difficult due the differences in testing methodology and subjects’ characteristics [9,20]. In our study, FAT_min_ was found to be at 53.40 % of VO_2max_ (range 49.40 to 62.52), which is lower than previous studies performed on athletic and sedentary men [3,7,8]. However, sedentary and adipose subjects cease to oxidize fat at lower exercise intensities due to weak aerobic and cardiorespiratory fitness [3,21]. Therefore, the less fit the subject, the earlier anaerobic metabolism will occur during an exercise, providing explanation for our observation [21].

Absolute MFO at FAT_max_ in this study was 0.72 g·min^−1^. These results are in line with previous studies where values from 0.30 to 0.70 g·min^−1^ have been reported [4,25]. Factors such as participants’ age, body composition, daytime, diet and testing methodology can affect substrates utilization and thus should be considered [25,26]. Compared to previous studies, several components induced higher lipolysis in our study. Previous studies had their subjects tested after high CHO meal either in overnight fast or in a post-prandial state, where our participants were tested after an overnight fast and after consuming LCHPF dinner [17,18]. Running involves larger muscle activity, hence causes higher lipolysis compared to cycling [27]. Consequently, the relative and absolute rate of fat oxidation in our study, compared to previous studies, was calculated using different stoichiometric equations, making direct comparisons hindered [20,25]. Determining which stoichiometric equation is the most suitable requires further examination.

The second aim of our study was to examine the existence of correlation between FAT_max_ and AeT and between FAT_min_ and AnT in men with obesity. Our results described a high correlation between the VO_2_ at AeT and at FAT_max_ (*r* = 0.89, *p* < 0.01) contrary to previous studies. Study by Bircher et al. [9] found low correlation (*r* = 0.32, *p* < 0.05) between AeT and FAT_max_ during cycle ergometry while using LT to determine thresholds. Different to our study, they used additional cycle test with 20 min stages and only three exercise intensities (55, 65, 75% VO_2max_) to determine FAT_max_. Fat oxidation was calculated only during last 5 min of each stage. Same group used identical protocol with more exercise intensities (25, 35, 45, 55, 65 and 75 % VO_2max_), determined threshold using VT and reported slightly higher correlation (*r* = 0.38, *p* < 0.05) [14]. Haufe et al. [4] found modest correlation (*r* = 0.55, *p* < 0.05) between LT and FAT_max_ while performing cycle test with 2 min stages to asses both VO_2max_ and FAT_max_.

Existence of high correlation between FAT_min_ and AnT found in our study (*r* = 0.99, *p* < 0. 01) is in line with previous studies performed on athletes and sedentary subjects where short test stages and VT were used [7,8,15]. It is believed that an increased lactate production in specific muscles and intra-muscles hypoxic conditions at intensities at/above AnT impedes lipolysis directly contributing to this phenomenon [24,28]. However, this occupancy requires further examination.

Several studies questioned the precision of substrates utilisation measurements at high exercise intensities during a non-steady state conditions which could possibly contribute to errors in correlation assessment [7,20]. Under these conditions indirect calorimetry is not necessarily representative of muscle substrates utilization since RER can be affected by changes in O_2_ and CO_2_. Still, the agreement exists how assessment is objective up to intensities of 85 % VO_2max_. The study by Jeukendrup et al. [20] advocates a possible shift in the acid-base balance during exercise intensities above 85 % VO_2max_. The high hydron ion will be buffered and ultimately, excessive CO_2_ exhaled through hyper-increase in BF_max_. This ventilator compensation will increase VCO_2_ and consequently result in overestimating CHO and underestimating fat oxidation. Considering how FAT_max_/AeT and FAT_min_/AnT in our study occurred at 42.80 % VO_2max_ and 53.40 % VO_2max_, respectively, we consider the measured values to be valid. Furthermore, in the attempt to maintain a stable bicarbonate pool, all our participants maintained a normal BF_max_ thus possibly diminishing any additional bias in the substrates utilization measurement [19].

When summarizing data from previous studies, low-to-modest correlations between FAT_max_ and AeT have been observed in adult sedentary men with obesity during cycle ergometry compared to our observations on a treadmill. Considering how studies demonstrated no difference in FAT_max_ location when cycle and treadmill ergometry are compared, physical activity type apparently will not influence the correlation [27]. Hence, we consider a testing methodology to be a main causer for previous observations; using prolonged test stages and reduced number of exercise intensities to determine FAT_max_ accompanied with usage of lactate could directly influence the accuracy in the identification of correlation by displacing FAT_max_ and metabolic thresholds [4,29].

Recent studies endorsed usage of short stage test as a gold standard to determine the VO_2max_ and the exercise intensity at which FAT_max_ occurs whereas tests with prolonged stages may not be so effective [6,7,30]. Furthermore, short-stage protocols have been validated to be a more suitable method when examining the correspondence between exercise intensity and fat oxidation in obese individuals since they provide linear VO_2max_ increase and an adequate time to utilisation steady state, increasing both metabolic thresholds and FAT_max_ determination precision [16,30,31]. Nevertheless, previous studies examining correlation in obese subjects did not use them. Considering the variety of techniques employed in determining the FAT_max_, caution should be taken as it seems that using prolonged stages tests provides inadequate response during FAT_max_ and VO_2max_ determination directly influencing correlation [6,30].

Apart that, studies demonstrated how long stage durations are rather time consuming and have an influence on lactate concentrations at the intensity eliciting the FAT_max_, additionally contributing to correlation assessment bias [30–32]. Contrary to LT, using VT demonstrated to be a better index of estimating metabolic thresholds in diverse subjects [2,12,28]. Moreover, the use of LT requires the continuous monitoring of lactate levels that is not an easy practice in subjects with obesity making VT a more practical mean [1–3]. Although LT and VT are causally related, they are controlled by different mechanisms which could direct affect correlation [11,28]. Accordingly, VT may be a more precise method when correlating metabolic thresholds with fat oxidation since the highest rate of oxygen extraction by the working tissue corresponds more accurately to the point of optimal ventilatory efficiency where levels of fat oxidation are maximal [11,28,29].

## Conclusion

The main findings of this study are high correlations between AeT & FAT_max_ and AnT & FAT_min_ in adult sedentary men with obesity while using a short stage treadmill GXT and ventilatory gases to detect metabolic thresholds. The presence of a low correlation between metabolic thresholds and fat oxidation in previous studies does not necessary mean lack of relationships between parameters, but rather the existence of different variables influencing the correlation. Hence, future studies should consider each of these variables and identify ways to reduce their effect. Our results suggest how metabolic thresholds and associated HR can be used as exercise intensity markers concerning maximal and minimal fat oxidation rate in adult sedentary men with obesity. These findings provide relevant and applicable information for further exercise prescriptions and recommendations.

